# Phage defence system abundances vary across environments and increase with viral density

**DOI:** 10.1101/2025.01.16.633327

**Authors:** Sean Meaden, Edze Westra, Peter Fineran

## Abstract

The defence systems bacteria use to protect themselves from their viruses are mechanistically and genetically diverse. Yet the ecological conditions that predict when defences are selected for remain unclear, as substantial variation in defence prevalence has been reported. Experimental work in simple communities suggests ecological factors can determine when specific defence systems are most beneficial, but applying these findings to complex communities has been challenging. Here, we use a comprehensive and environmentally balanced collection of metagenomes to survey the defence landscape across complex microbial communities. We also assess the association between the viral community and the prevalence of defence systems. We identify strong environmental effects in predicting overall defence abundance, with animal-host-associated environments and hot environments harbouring more defences overall. We also find a positive correlation between the density and diversity of viruses in the community and the abundance of defence systems. This study provides insights into the ecological factors that influence the composition and distribution of bacterial defence systems in complex microbial environments and outlines future directions for the study of defence system ecology.

## Introduction

The immense diversity and complexity of the global virome has resulted in a multitude of ways that bacteria can defend themselves against viral infections. These defences span a range of mechanisms, including restriction modification (RM) and CRISPR systems, which recognise and cleave infecting viral genomes, while abortive infection systems often trigger degradation of essential molecules of the host cell, preventing further spread of viral particles [1]. In recent years, many more types of defence systems have been identified, expanding the known ‘defensome’ [2] and this diversity of defences and virus-encoded counter defences [3] suggest ongoing co-evolution and ecological significance. Many of these defence systems frequently co-occur within the same genome [4] with some combinations providing additional levels of defence through both additive and synergistic interactions [5–7]. In general, defence systems exhibit broad genetic and mechanistic diversity, encompassing those that degrade invading nucleic acids, such as RM and CRISPR, those that trigger cell death or dormancy, such as abortive infection (Abi) systems like toxIN, or type-III CRISPR and CBASS, and those that exhibit numerous other mechanisms (reviewed in [8]). These diverse mechanisms may also be favoured under specific ecological scenarios (reviewed in [9]), for example systems that protect neighbouring cells may be more beneficial when environmental spatial structure is high [10]. However, linking these defence systems to environmental factors in complex environments is challenging and the ecological drivers that shape their distributions in natural environments are less well understood. Finally, defence systems are frequently carried by mobile genetic elements (MGEs), allowing rapid mobilisation into new hosts and facilitating competition between MGEs for shared hosts (reviewed in [11]).

Both theory and experiments have revealed potential drivers of the evolutionary ecology of defence systems (reviewed in [9]), but we lack a synthesis of this knowledge and many open questions remain in complex microbial communities (reviewed in [12]). Abiotic factors generally determine microbial community composition (e.g. pH or salinity, [13]) and these vary substantially across environments, as shown by the Earth Microbiome Project (EMPO)[13]. In turn, biotic factors such as virus:microbe ratios, population sizes and interaction rates also vary across environments and these biotic factors are likely to shape the composition of defence systems present [12]. For example, high relatedness between neighbouring bacteria has been shown to determine when abortive infection is a successful strategy [10], while the benefits of CRISPR systems outweigh receptor-based resistance when microbial biodiversity is high [14], whereas defence can be selected against in the presence of beneficial antibiotic resistance encoding plasmids [15]. As defence systems can be readily gained and lost, or inactivated, from bacterial genomes [15–22], their distributions may be optimised by ecological factors.

Genomic surveys of defence systems across bacterial and archaeal genomes have revealed some of these drivers of defence system composition. Multiple studies have found strong effects of genome size on the abundance of defences [23,24]. Furthermore, by linking genomes to predicted traits, additional drivers can be inferred, such as aerobicity [25] and temperature [26] for CRISPR systems, temperature for RM systems and fast growth rates for overall defence abundance [27]. Expanding this approach to include genomes assembled from metagenomic data has shown that genomes from the gut environment carry substantially more defence systems than genomes from soils and the oceans, respectively [2], and that plant-associated bacteria have fewer defence systems than non-associated close relatives [28].

An additional factor predicted to shape the abundance and type of defence systems present in an environment is the density of viruses. Higher density would likely lead to more frequent infections and, in turn, stronger selection for defence systems. We previously identified that the abundance of CRISPR defence systems is both positively correlated with the abundance of viruses, and varies widely across different microbial environments, with host associated environments carrying more CRISPR than free-living environments [29]. Here, we collected a diverse range of metagenome assemblies from a public sequence data repository (MGnify, European Nucleotide Archive, [30]) and mined these assemblies for both defence systems and viral sequences. We then used coverage information as a proxy for the relative abundance of defence systems and viruses. We describe the distribution of defence systems across environments, variation in the total amount of defence systems and link defence abundance to the abundance, or density, of viruses in each sample.

## Methods

### Data curation

We collected a wide-ranging and standardised collection of metagenomes that had been processed using consistent methods. The MGnify database (European Nucleotide Archive, ENA) was accessed via the API using the MGnifyR R package (https://github.com/EBI-Metagenomics/MGnifyR) on 14/06/2024. A search was conducted for all MGnify samples labelled as ‘assembly’ under the ‘experiment-type’ field and filtered to retain those processed using the MGnify pipeline version 5.0. Metadata from the resulting 34,799 assemblies was collected with the getMetadata function from MGnifyR. Samples were then selected from categories in the ‘biome_string’ field that had > 99 samples per group with a requirement that only Illumina sequence data was included. The locations of the assemblies within the ENA database were located with searchFile function of MGnifyR and the resulting URLs downloaded via a curl command on the University of York’s HPC server. The associated fastq data files for each assembly were downloaded using enaBrowserTools [31] and subsampled to 1 million reads per sample using seqtk [32]. Samples with fewer than 1 million reads were excluded from downstream analyses.

### Metagenome community composition and diversity

From the subsampled 1 million reads from each sample, 500,000 were taxonomically profiled using Kraken2 with the Kraken2 standard database [33]. Taxonomic groups that made up less than 0.01% of the relative abundance were removed. Diversity at the Genus level was calculated and PCoA clustering performed using the ‘vegan’ R package [34].

### Defence system search

Contigs were first annotated for coding sequences using prodigal (default settings, translation table 11) [35]. Comparison on a subset of samples found that when ‘meta’ mode was used the results were almost identical. Assemblies shorter than 100kbp were excluded. Contigs were searched for defence systems using PADLOC ([36], version 2.0) with the PADLOC database (version 2.0). Defences labelled as ‘DMS_other’, ‘dXTPase’, ‘PDC’, ‘HEC’ and ‘VSPR’ were discarded as these are either non-defence, not experimentally verified or unpublished predicted defence systems at the time of analysis, although some HEC systems have been subsequently verified [37]. Assemblies were searched for CRISPR arrays with metaCRT using default parameters [38].

### Viral sequence search

Assemblies were searched for viral sequences using the geNomad pipeline [39]. Sequences shorter than 1 kb and those lacking any ‘hallmark’ (as classified by geNomad) viral genes were discarded. Sequences were classified as chromosomal, plasmid or phage based on the maximum score ascribed by geNomad. The genomic locations of defences were identified using the geNomad predictions for chromosome, phage, plasmid or prophage again using the maximum score.

### Abundance estimation

The associated sequencing reads were mapped to the assemblies with bwa ([40], version 0.7.17) and processed with samtools ([41], version 1.9). The resulting alignment files (BAM format) were used to calculate coverage and read recruitment values for all contigs in each assembly using coverM with the ‘contig’ command (version 0.7.0) and the ‘metabat’ and ‘count’ methods respectively. The contigs identified from the defence system and viral sequence searches were then extracted from the coverM results tables and used for downstream analysis. These per-contig abundance files were then merged with the PADLOC results with 1 count value recorded per defence system-type per contig. In cases where one contig carried multiple defence systems, the count value was recorded for each defence type. Notably, of all the contigs identified as carrying a defence system, >95% carried a single defence system type, likely due to the highly fragmented nature of the assemblies (Fig. S1). The total defence abundance per sample was calculated by the sum of reads mapping to contigs carrying each defence system. We opted here to focus on measuring the abundance of defence systems and therefore count multiple unique defence systems on a single contig independently, effectively ‘double counting’ contig read recruitment if it carried multiple unique defences. In practice, this represented <5% of all contigs due to the fragmented nature of the assemblies (4% carrying 2 systems, 0.4% carrying 3 systems and 0.5% carrying 4 or more systems). Count data was also obtained using the same mapping-based approach for those contigs predicted to be of viral origin: abundance tables were extracted from the coverM results tables based on the geNomad predictions and collated into a master file containing the results from all samples.

### Sequencing effort and eukaryotic contamination controls

We assessed the effect of sequencing depth by first collecting the associated fastq data and counting the number of reads. We also assessed eukaryotic (human) DNA contamination, predicted from the Kraken2 analysis, and found higher levels of eukaryotic DNA associated with human-associated samples (Fig. S2). When analysis was restricted to samples with < 50% of classified reads being of eukaryotic origin, and with the inclusion of assembly N50 and original sequencing depth, the results were qualitatively the same. There remained a strong environmental effect on defence abundance and a consistent correlation between viral abundance and defence abundance.

### Taxonomic identification of plant-derived defence systems

We observed a high number of argonaute and Tiamat systems in the plant-associated samples. To assess the origin of these defence systems we extracted the corresponding contigs, on which these defences were located, and assigned a taxonomic classification using the MMSeqs2 taxonomy pipeline against the NCBI non-redundant (nr) nucleotide database (downloaded 25/08/2024). Contig classifications based on the last common ancestor (LCA) were visualised in R.

### Statistical analysis

The effect of environment on total defence abundance was assessed using a GLM with a ‘quasipoisson’ error structure. A measure of assembly fragmentation (N50 value) was included in the model, as preliminary analysis found a significant relationship between N50 value and defence abundance. Significance was assessed with ANOVA by comparison against a null model with the environment term removed.

The effect of environment of defence system composition was assessed by first converting the results to a sample by defence system abundance matrix. Permutational ANOVA was then applied using the ‘adonis2’ function from the ‘vegan’ R package. This function converts the abundance matrix to Bray-Curtis distances of dissimilarity between samples and applies a permutational ANOVA. 999 permutations were used. Ordinations were conducted using the ‘NMDS’ and ‘pco’ functions in ‘vegan’ based on Bray-Curtis dissimilarity values.

Correlations between defence abundance and viral abundance were assessed using a linear model including N50 as a covariate and a gaussian error structure. Defence abundance and viral abundance counts were log10 transformed to improve model fit and model residuals were assessed visually. Significance was assessed by ANOVA against a null model with viral abundance removed. We repeated the above statistical tests on a restricted dataset that had the following criteria: all samples had < 50% of reads classified as eukaryotic and both the assembly N50 value and the read count of the original samples included in the model. This aimed to account for assembly fragmentation and the sequencing depth, based on the assumption that all the data were used to generate each the assemblies.

## Results

### Defence abundance, diversity and composition varies across environmental categories

The discovery of novel defence systems continues at pace, but our knowledge of the ultimate drivers of their evolution and ecology is lacking. Here, we collated an ecologically diverse collection of 1075 metagenomes from 12 environments. We then surveyed the defence repertoire and abundances using a homologue-search based tool [36] to assess which environments carry the most defences and the composition of defence systems in those environments. We found that the total abundance of defence carrying contigs significantly varied among environments (F_11,1037_ = 46.0, p < 0.0001), suggesting that environmental conditions strongly affect when defence systems are optimal. Notably, animal-host associated environments (urethra, gut, vagina, oral and skin) were highest in overall defence abundance, with hot environments similarly high in defence abundance (Fig. 1).

**Fig. 1.**
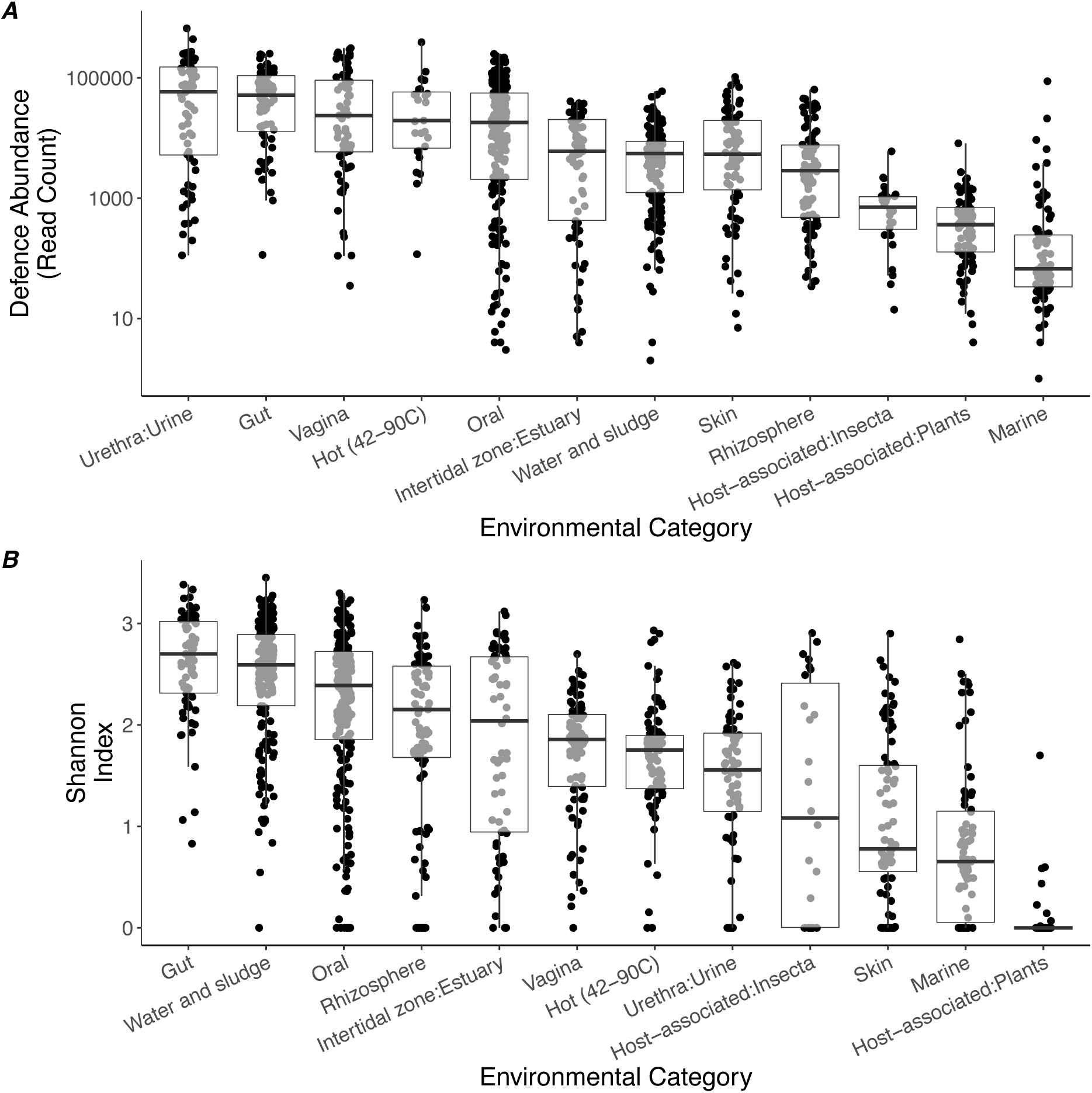
Defence abundance and diversity varies across microbial environments. A) Points represent individual metagenomes grouped by biome metadata categories provided from the European Nucleotide Archive. Defence systems were identified from metagenomic contigs using PADLOC [36], a standardised subsample of 1 million reads were mapped to each assembly and the counts of contigs carrying defence systems were summed to count total defence per sample (>95% of contigs had a single defence system). Boxplots show median values and first and third quartiles. B) Boxplots showing defence diversity (Shannon Index) for each environment. Points represent individual metagenomic assemblies. Shannon’s diversity index was calculated from a subsample of 1 million reads mapped to contigs carrying defence systems.

We then calculated the diversity of defences (Shannon’s index) for each sample, which also varied significantly with environment (F_11,1064_ = 75.2, p < 0.0001, Fig. 1). Overall, defence diversity was correlated with defence abundance (F_1,987_ = 719.9, p < 0.0001); however, there were minor differences in the ranking of the environmental categories. Water and sludge, estuary and soil (rhizosphere) environments had similarly high defence diversity to the host-associated samples, despite lower defence abundances. We also observed substantial variation in the abundance and distribution of specific defences, with a notably sparse distribution, meaning that most defences were rare or absent in the majority of samples (Fig. S3). To assess the role of the environment in shaping the defence composition we applied a permutational ANOVA to a Bray-Curtis matrix of dissimilarity between samples using environment as a fixed effect. We found a significant effect of environment (F_11,1133_ = 23.0, p < 0.001) with an R^2^ value of 0.18. Despite this significant effect, inspection of NMDS ordinations showed substantial overlap between groups (Fig. 2A), preventing accurate prediction of defence composition from environmental information alone. Importantly, taxonomic profiling of the same samples did show strong clustering by environment (Fig. 2B) and we observe a substantial mismatch between overall community diversity and defence diversity (Fig. S4 and Fig. 1B). In agreement with the pan-immune hypothesis of rapid gain and loss of defences [42], these results suggest that differences in defence composition are not driven solely by bacterial taxonomic effects.

**Fig. 2.**
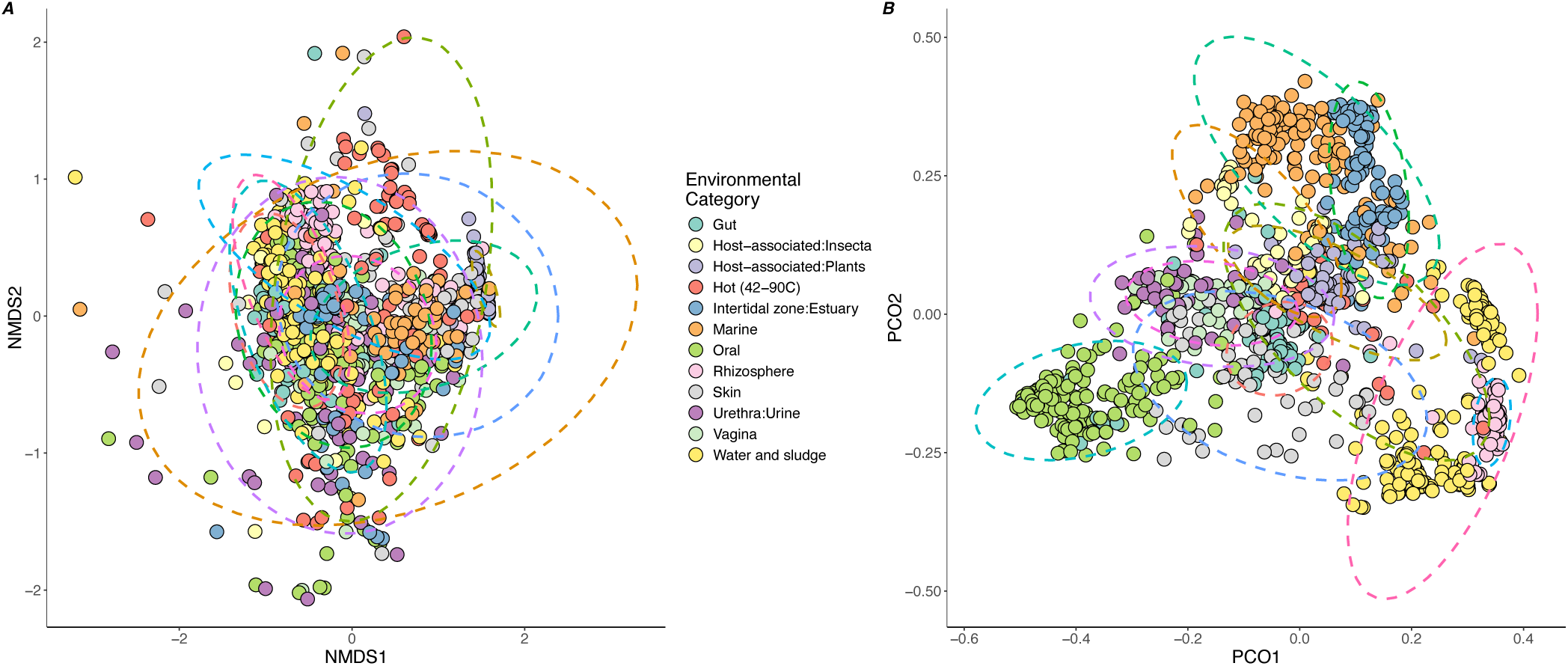
Defence and taxonomic composition variation between environments. Ordinations of metagenomic samples based on similarity in defence composition (A) or taxonomic similarity (B). Groups that cluster together share more defence systems or prokaryotic taxa respectively. Points represent individual metagenomic assemblies, ellipses represent 95% confidence intervals for a multivariate t-distribution and colours show the environment sampled. A) NMDS analysis was performed on a defence system abundance table constructed from a subsample of 1 million reads mapped to contigs carrying defence systems. B) A subsample of 500,000 reads were classified with Kraken2 and the frequency of each genus collected. Both ordinations use Bray-Curtis dissimilarity scores calculated from their respective abundance tables.

### Defence abundance correlates with viral abundance across environments

In addition to abiotic factors, biotic factors (and in particular the viral community composition), are likely to influence the abundance of phage defence systems. To assess this, we first estimated the abundance of viruses in the sample based on coverage of viral contigs from a standardized subset of reads for each sample (1 million). These estimations therefore do not represent absolute viral abundances which likely vary substantially between environments, but measure viral abundance relative to microbial DNA sequences as the majority of reads are of bacterial origin. We found a significant correlation between overall defence abundance and viral abundance (F_1,997_ = 115, p < 0.0001, Fig. 3), suggesting that the density of viruses is a strong selective force for phage defence systems. As an additional test, we restricted the analysis to just those viruses annotated as Caudoviricites as a way of excluding effects caused by non-phage environmental viruses. In this case we also observed a significant positive correlation with overall defence abundance and Caudoviricites abundance (F_1, 978_ = 135.0, p < 0.0001, Fig. 3). Unsurprisingly, viral diversity and viral abundances were strongly correlated (Pearson correlation coefficient: 0.39, p < 0.0001). We refer herein to viral abundance but note that the accompanying viral diversity may also be contributing to the observed effects. We also identified the predicted genomic location of each defence system to assess the proportions of defences carried on MGEs. We found 54% of defences were located on chromosomal contigs, 32% on plasmid contigs, 13% on viral contigs and 0.3% on integrated prophages (Fig. S5). We note that many of the viral contigs likely represent fragments of prophages and the low number of integrated prophages results from the scarcity of fully intact prophage genomes complete with chromosomal flanks due to fragmented metagenomic assemblies.

**Figure 3.**
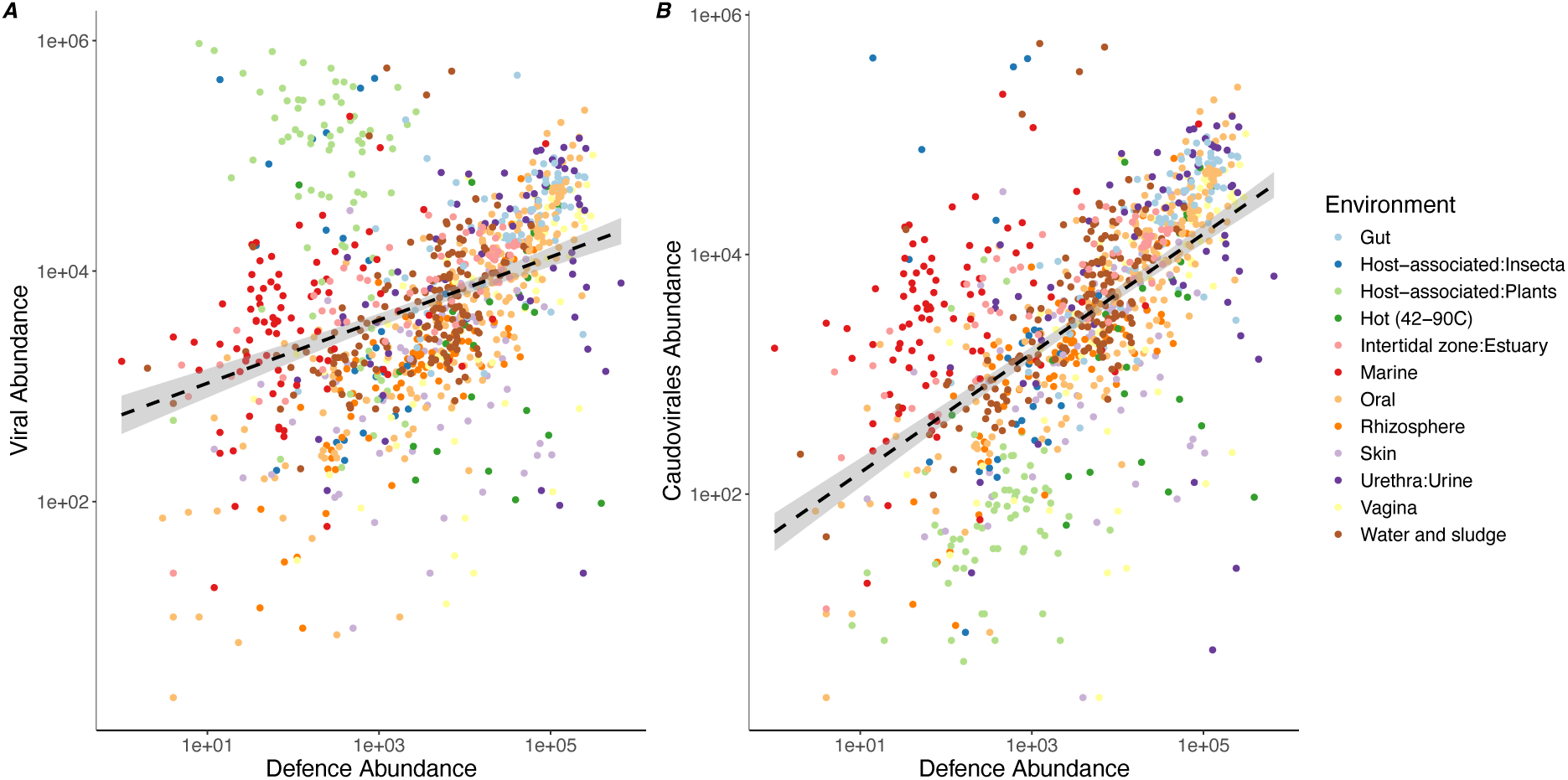
Defence abundance correlates with viral abundance. Metagenomic assemblies were mined for phage defence systems and viral sequences. Coverage values were collected from a subsampled of 1 million reads per sample. Viral abundance represents the sum of all read counts for virally classified contigs; defence system abundance represents the sum of reads mapping to contigs carrying defence systems. The dashed lines represent linear models and the shaded area 95% confidence intervals. Panel A) shows the sum of all viral contig read counts while B) shows a subset of counts restricted to contigs annotated as the dsDNA tailed phage group Caudoviricetes.

### CRISPR array abundance correlates with viral abundance across environments

Our previous work assessed the abundance of CRISPR defence systems across environments. We aimed to use a similar approach across the wider pool of samples used here, again focusing on CRISPR arrays rather than effector proteins. By focusing on CRISPR arrays we aimed to mitigate the difficulties of detecting multi-gene systems in fragmented, short contig, metagenomic data. As before, we found a positive correlation between CRISPR array abundance and viral abundance (F_1,911_ = 4.33, p = 0.04) and a similar relationship when we restricted the analysis to Caudoviricites abundances (F_1,896_ = 9.56, p < 0.01). We also assessed the origins of the arrays and found that similar numbers of arrays were predicted to be located on chromosomes (40.2%) and viral contigs (44.7%), which include prophages, with fewer predicted on plasmids (15.1%). When we repeated the correlations between CRISPR abundance and viral abundance for these subsets of the data, we found significant positive correlations between CRISPR abundance and viral abundance for the chromosomal (F_1,838_ = 208.0, p < 0.0001) and plasmid (F_1,697_ = 93.7, p < 0.0001) derived array abundances, but not virally derived arrays (F_1,814_ = 0.94, p = 0.99). Taken together these results suggest that the selective forces that determine when CRISPR is beneficial may differ between bacteria and plasmids vs. phages, potentially due to stronger selection for streamlined genomes in viruses.

### Reduced defence system prevalence in plant-associated environments

While the abundance of defences generally correlated with viral abundance across environments, the plant metagenomes used in this study were obvious outliers, harbouring a highly reduced number of defence systems and elevated viral abundance (Fig. 3A). Plant microbiome samples typically suffer from high levels of plant DNA contamination due to sampling techniques [43]; however, recent work has also found phage defence systems to be underrepresented in plant environments [28]. The defence systems most abundant in the plant samples in our dataset were PD-T4, argonautes and Tiamat. Argonautes are well characterised plant immune effectors in RNA silencing immunity [44] and are therefore unsurprising to be prevalent, if the metagenome is largely plant-host derived. To assess the taxonomic origins of argonaute and Tiamat systems we extracted the contigs containing these systems and classified them using the NCBI non-redundant nucleotide database. For the plant associated metagenomes, 86% of argonaute and 100% of Tiamat systems were located on plant genome sequences (Fig. S6). These observations of plant-derived sequences are supported by the presence of many viral sequences annotated as RNA viruses, despite the data being from metagenomic data, which can occur in the data as endogenous retroviruses integrated into the plant genome. Despite these technical aspects of plant microbiome sampling, further work must assess the biological reasons for an underrepresentation of phage defence systems in plant environments [28].

## Discussion

Previous work has identified variation in both the frequency and types of phage defence systems found across different natural environments [2]. We have previously shown that the abundance of a specific defence system, CRISPR-Cas, is strongly correlated with the relative abundance of viruses present in the environment [29]. Here, we conduct a survey of the defence systems present in a broad range of environments using publicly available metagenomic data. We found a strong correlation with the total abundance of defence systems and viral abundance, consistent with the notion that the viral community is a strong selective force for the acquisition and retention of phage defence systems.

In agreement with other work, we found that gut samples harboured a greater abundance of defence systems than soil or marine environments respectively [2]. Notably, five out of the top six environments for viral abundance are human-host associated, with the exception being samples derived from environments 42°C or higher (such as hot-spring thermal environments). Hot-spring environments are suggested to be a hotspot for virus-defence systems due to higher costs associated with mutations, via reduced protein stability, in turn reducing viral diversity and the potential for defence evasion [45]. By contrast, bacteria and archaea from mesophilic environments may be more able to tolerate a wider range of mutations, requiring more robust resistance mechanisms, such as mutation of surface receptors and subsequent phage resistance. Indeed, recent work suggests surface receptor variation can be a stronger predictor of successful phage infection than intracellular defence systems [46]. Our results are consistent with the notion of human-host environments providing a resource rich environment for microbes, in turn hosting a greater density of viruses and selection for defence systems.

Interestingly, marine environments appear to be particularly depleted in defence systems. Although the marine environment is predicted to have a high daily viral lysis, which is consistent with strong selection for defence, the virus-to-microbe ratio (VMR) is low [12]. Further, infection assays of culturable marine microbes found low predation pressures likely due to low encounter rates [47]. In addition, the dominant marine clade, SAR11, has undergone genome reduction [48], presumably reducing the capacity for carrying diverse intracellular defence systems. The low overall density of defences we observed is consistent with relatively weak selection from phages and the typically low nutrient conditions may make the carriage of defence systems costly compared to selection for reduced genome sizes. In support of this conclusion, analysis of cyanobacteria and their phages found far more defence systems in freshwater genomes, which typically have higher nutrient availability, than those from marine environments [49]. In contrast to human host-associated samples, insect associated samples also carried far fewer defences. It has been observed previously that insect-associated bacterial genomes have few or no defences [23,50] and is likely either due to the general genome reduction processes that occur in intracellular insect symbionts [51] or reduced phage predation in the endosymbiotic environment (discussed in [52]).

Plant-associated samples also carried far fewer phage defence systems than human host-associated samples. The phyllosphere is typically low in carbon and nitrogen and relatively oligotrophic [53] again potentially increasing costs of phage defence system carriage. Recent work has found that plant-associated bacteria are depleted in defence systems relative to non-plant-associated relatives [28]. However, our results may also be partially due to technical artefacts of sampling plant tissue and the typically high levels of host contamination. Specifically, in our dataset we found argonautes to be the most abundant defence, which are common in plant genomes, functioning as RNAi effectors [44]. The viral community was also consistent with this conclusion as although Caudoviricites was the most frequently identified viral group, we found many groups of Riboviria. These are RNA viruses capable of integrating into plant host genomes as endogenous retroviruses. Surprisingly, along with argonautes we also identified a high frequency of the Tiamat and PD-T4-6 defence systems. When we identified the origins of the Tiamat and argonaute systems, these were almost exclusively from plant sequences in the plant samples, versus a wide range of bacteria in the other environmental samples (Fig. S6). Further work is needed to assess the reasons why the plant-associated metagenomes were so depleted in defence systems [28] and the extent of defence conservation across domains of life [54].

By mining existing metagenomic assemblies, our results may be skewed towards viruses that are enclosed within a cell, either as prophages or undergoing active replication [55], although some viral particles will be present, as bulk metagenomes typically containing the most abundant viral genomes [56]. Yet experimental work has shown that the extracellular viromic fraction in an environment can change quickly, both temporally and spatially [56]. In addition, we focus entirely on DNA viruses, but RNA viruses are common [57,58] albeit less well studied. Assessing patterns of defence prevalence in light of RNA virus abundances, and integrating spatial and temporal information will be important future work. We also cannot rule out some biases in our search strategy as most defence systems, including those derived from a wide range of non-model bacteria and archaea, have been functionally validated in a limited number of model organisms [59,60]. It is possible that some incompatibility between defence-system origin host and the taxa chosen as model organism creates biases in defence system discovery. We also note that our results are skewed towards smaller defence systems, with fewer core genes, due to the fragmented nature of typical metagenomic assemblies (Fig. S7). Longer, multi-gene defence systems are more likely to span multiple contigs in the assembly, and will therefore not be detected by defence identification tools, which rely on finding core genes and/or a minimum number of genes depending on the system. Finally, metagenomic assemblies are rarely exhaustive and likely represent the most abundant organisms in an environment; as such our defence system survey is representative of those that exist in the most abundant species and many others almost certainly exist in those environments at lower frequencies. Future efforts must focus on more contiguous assemblies derived from long-read sequencing that will be vital for more fine scale metagenomic analysis.

We found a consistent positive correlation between defence abundance and viral abundance; however, viruses and other MGEs are well-known to carry defence systems. Our analysis found up to 46% of defences predicted to be located on MGEs. Therefore, this high percentage of MGE associated defence systems may be driving the observed correlations. This leads to two possible interpretations: firstly, that defence accumulation on MGEs is a neutral process or ‘lottery’ effect, and will occur more frequently when MGE abundances are high, or secondly, that when MGE abundances are high there is greater competition between MGEs for susceptible hosts, requiring more defence systems to target competitors [11,61]. We suggest that experimental work in this area will yield valuable insights into natural microbial community dynamics of MGE competition and the interplay with defence systems.

Overall, we have identified wide variation in defence abundances across microbial environments and the density of viruses as a likely driver of selection for defence systems. Despite differences in defence abundance, there were minor differences in defence system composition across environments. This was surprising given the strong clustering of samples at the taxonomic level (Fig. 2) and suggests that while the environment predicts the overall abundance of defence, it does not strongly shape the defence composition. Clearly, further work that integrates community ecology and metagenomic analysis is needed to assess whether the accumulation of specific defences is a stochastic process or determined by other unmeasured parameters. We also predict that further work may identify the ultimate selective forces acting on individual defence systems. As ever more defence systems are discovered we anticipate future studies will focus on the individual ecology of system types and classes of defence [8] and anti-defence [3], potentially identifying environmental hotspots that would allow targeted search strategies for defence discovery.

## Data and Code Availability

All data used are publicly available from the MGnify database. A list of accession numbers is found in supplementary file 1. Code used for the analysis is available at github.com/s-meaden/mg_dfs.

## Acknowledgements and funding statement

SM was supported by funding from the Biotechnology and Biological Sciences Research Council [BB/X009793/1]. This work was further supported by the Biotechnology and Biological Sciences Research Council (sLoLa grant BB/X003051/1), a UK Research and Innovation grant under the UK Government’s Horizon Europe funding guarantee (EP/X030377/1), and the Philip Leverhulme Prize (PLP-2020-008) to E.R.W. PCF was supported by Bioprotection Aotearoa (Tertiary Education Commission, NZ) and by a James Cook Research Fellowship (RSNZ, Te Apārangi). For the purpose of open access, the author has applied a ‘Creative Commons Attribution (CC BY) licence to any Author Accepted Manuscript version arising from this submission.

## Supplemental Information

**Fig. S1.**
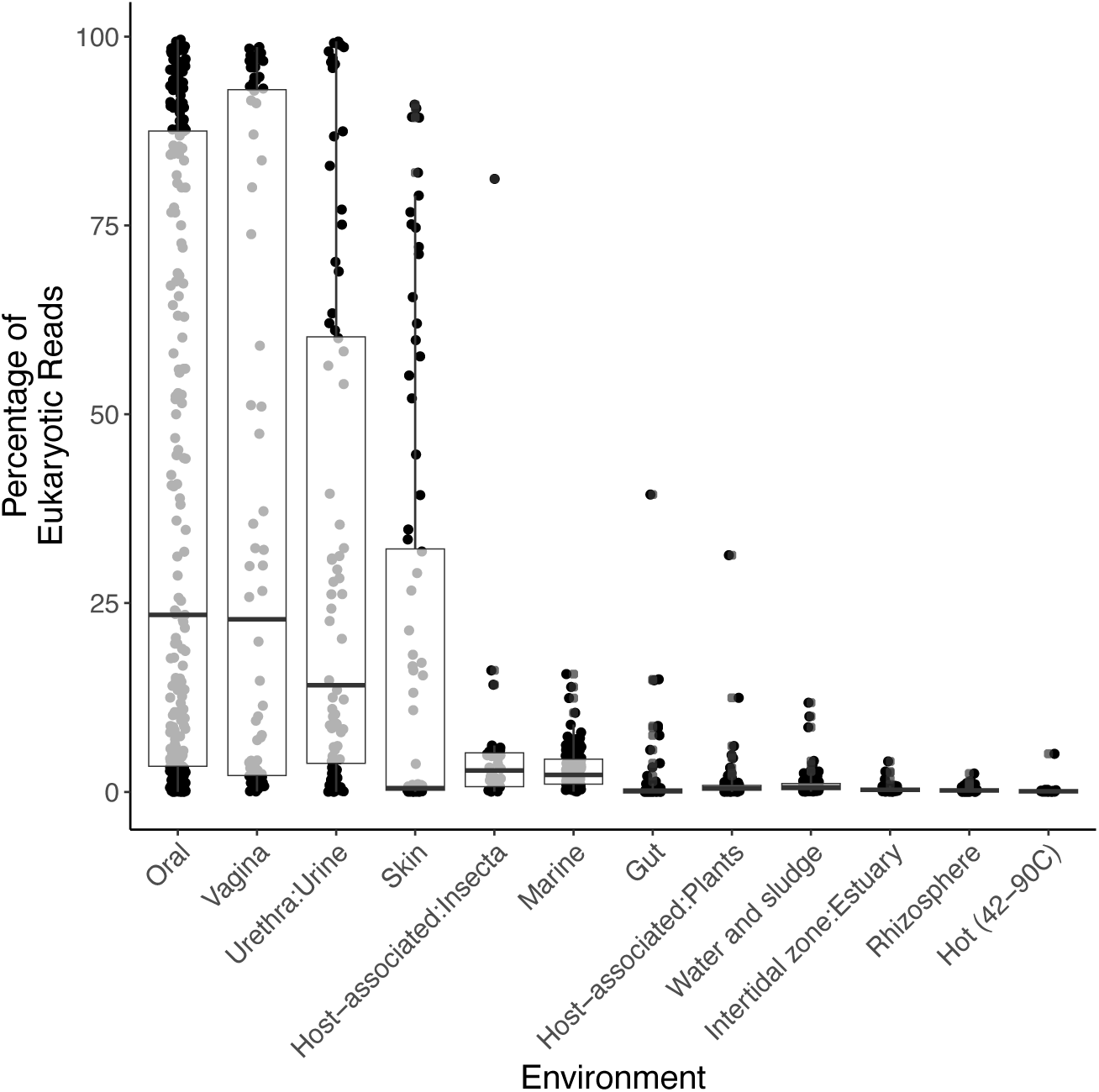
Percentage of reads classified as of human host origin. Boxplots showing the percentage of reads classified as eukaryotic from the Kraken2 classifier. Samples are grouped by environmental classification and ordered based on the median value. Each point represents an individual metagenome. Samples with >50% of reads being classed as eukaryotic were excluded from downstream analysis. Note that the standard Kraken2 database includes microbial genomes (bacterial, archaeal, plasmid, viral, UniVec vectors and the human reference genome) but not plant, fungal or protozoa.

**Fig. S2.**
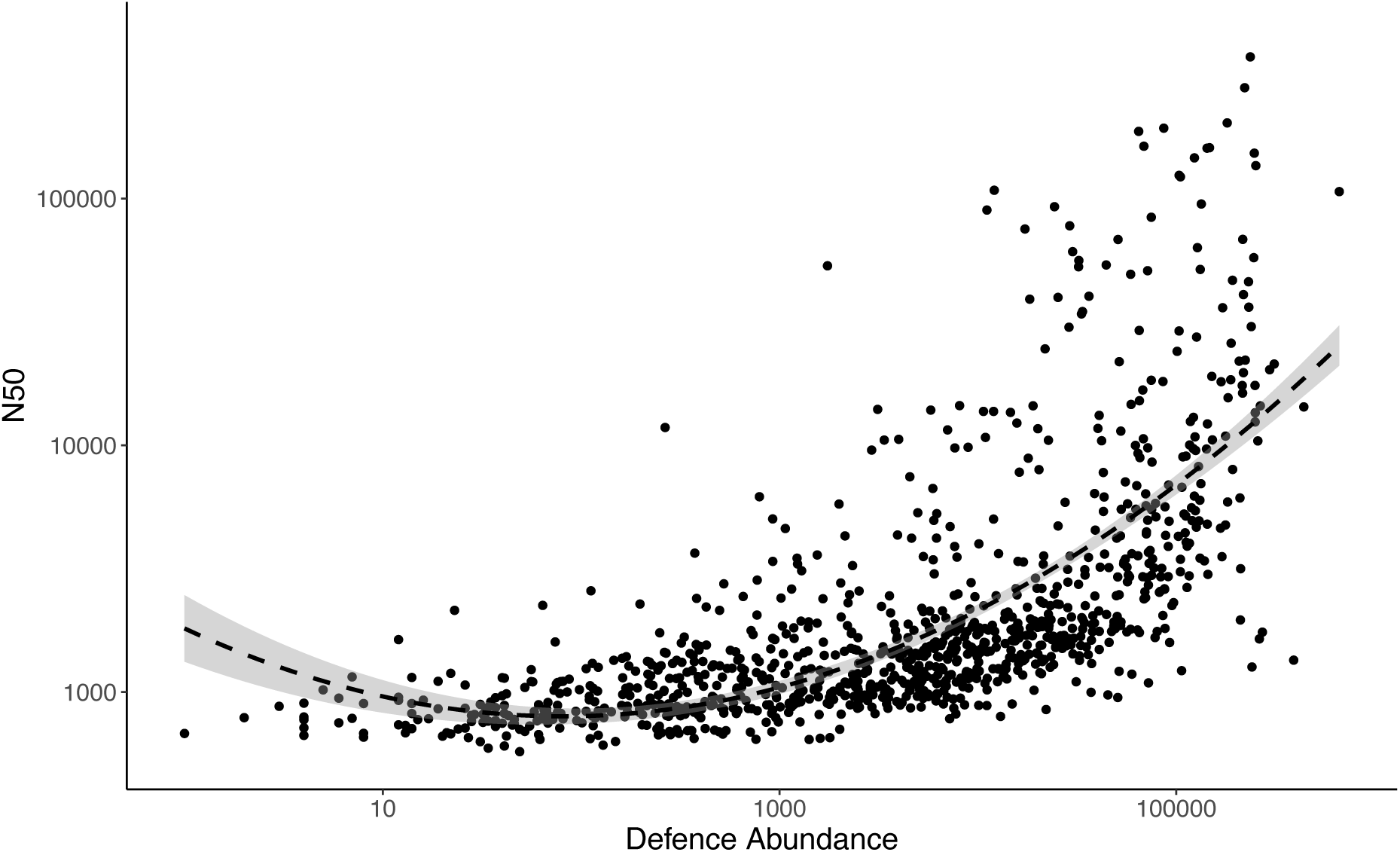
Metagenomes with more contiguous assemblies (greater N50 value) recover more defence systems. Correlation between the N50 value of each assembly and the count of reads that map to defence carrying contigs. N50 values represent the shortest contig length, when ranked by length, to be included to cover 50% of the genome and we use here as a proxy for assembly fragmentation. Points represent metagenomic assemblies and dashed line represents a polynomial linear model, with shaded areas showing 95% confidence intervals.

**Fig. S3.**
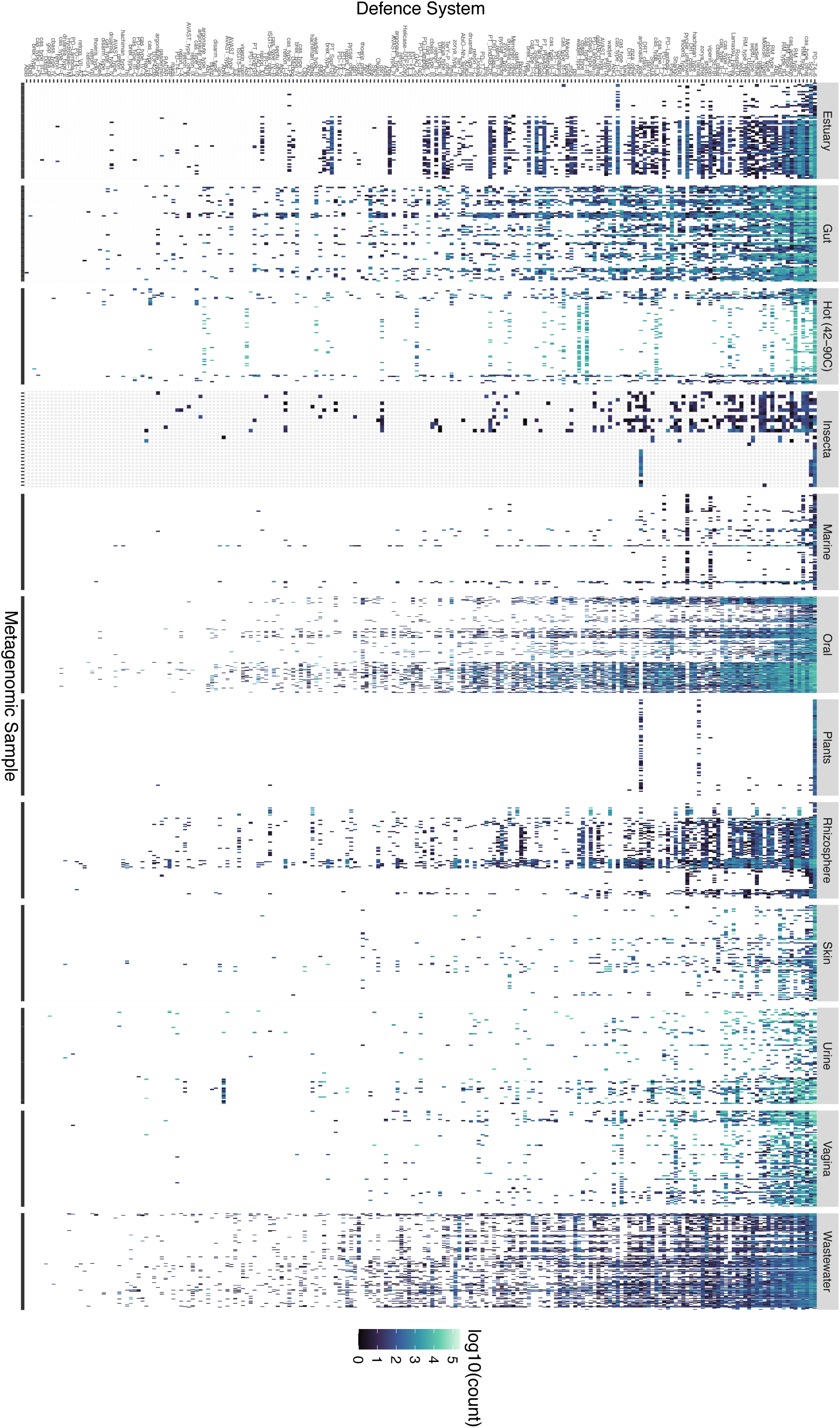
Variation in defence system distributions across microbial ecoystems. Individual columns refer to metagenomic assemblies, with groupings representing the microbial environments samples were collected from. Rows represent individual defence systems and grid colours represent the sum of counts of reads recruited to contigs carrying each defence system from based on a standardised subsample of 1 million reads per sample.

**Fig. S4.**
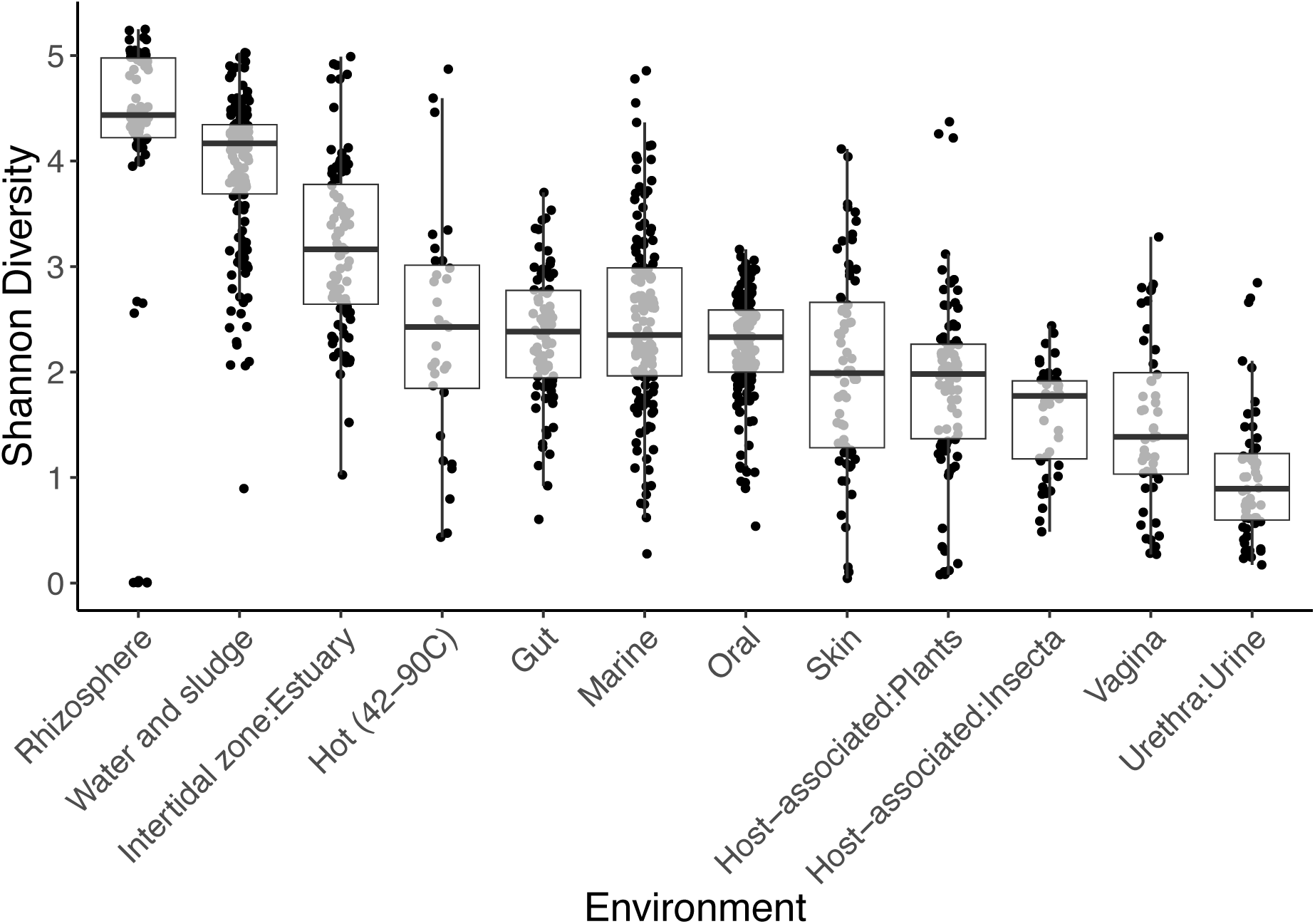
Microbial community diversity varies by environment. Boxplots of the Shannon Diversity values based on the genera identified in each sample from Kraken2 analysis. A subsample of 1 million reads were classified with Kraken2 and the frequency of each genus collected. Categories are ordered by the median value per environment. Note these counts include all identified genera, but largely reflect bacterial and archaeal diversity.

**Fig. S5.**
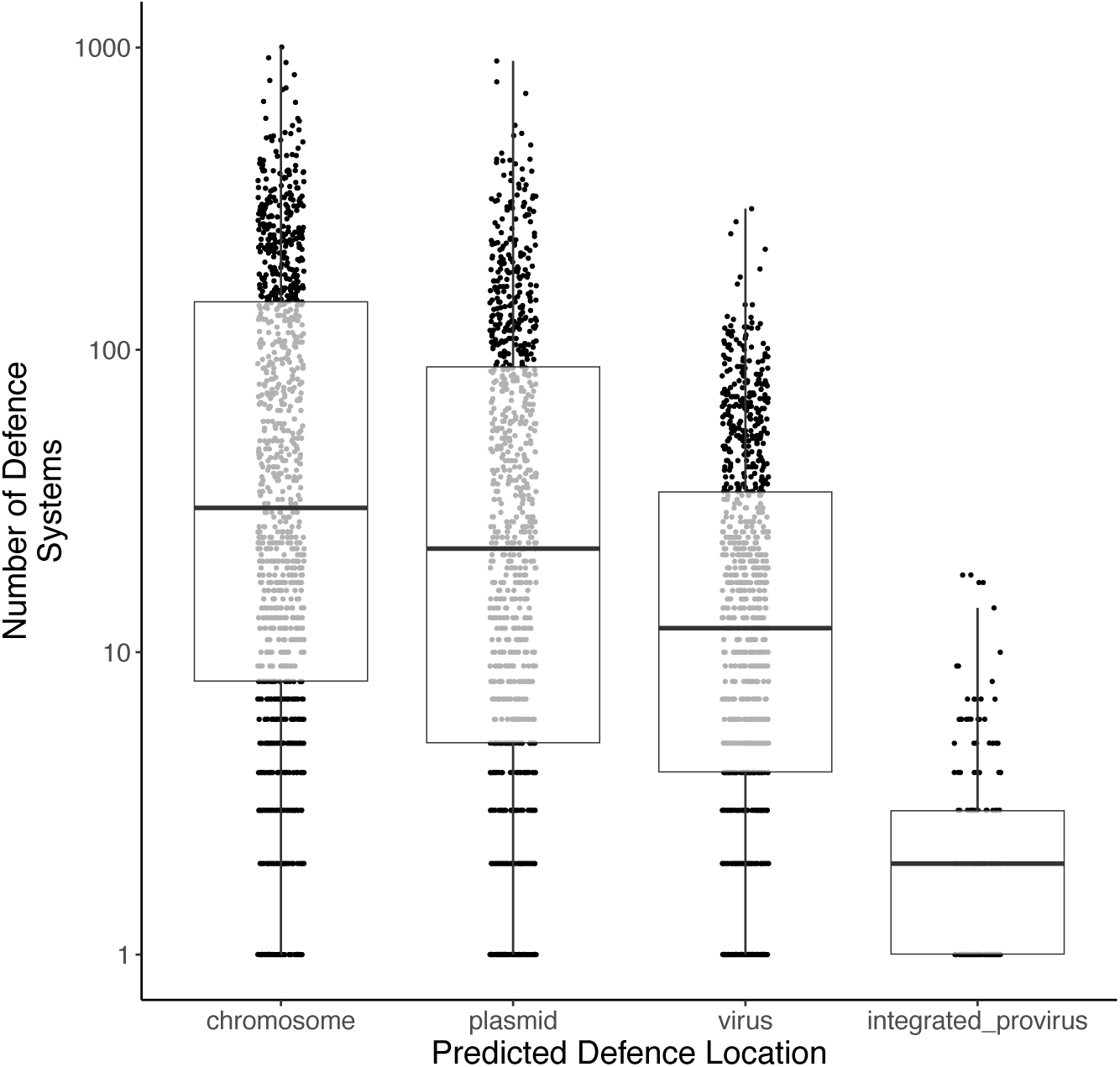
Predicted locations of defence systems. All defence system carrying contigs were classified with the geNomad pipeline and assigned as chromosomal, plasmid, viral or integrated proviral based on the maximum score assigned by geNomad. For those contigs carrying an an integrated provirus, the genomic coordinates of the provirus was used to assign the defence as proviral or chromosomally encoded. Note y-axis here represents the frequency of defence systems, not abundance. We also note that many of the viral contigs likely represent fragments of prophages and the low number of integrated prophages results from the scarcity of fully intact prophage genomes complete with chromosomal flanks due to fragmented metagenomic assemblies.

**Fig. S6.**
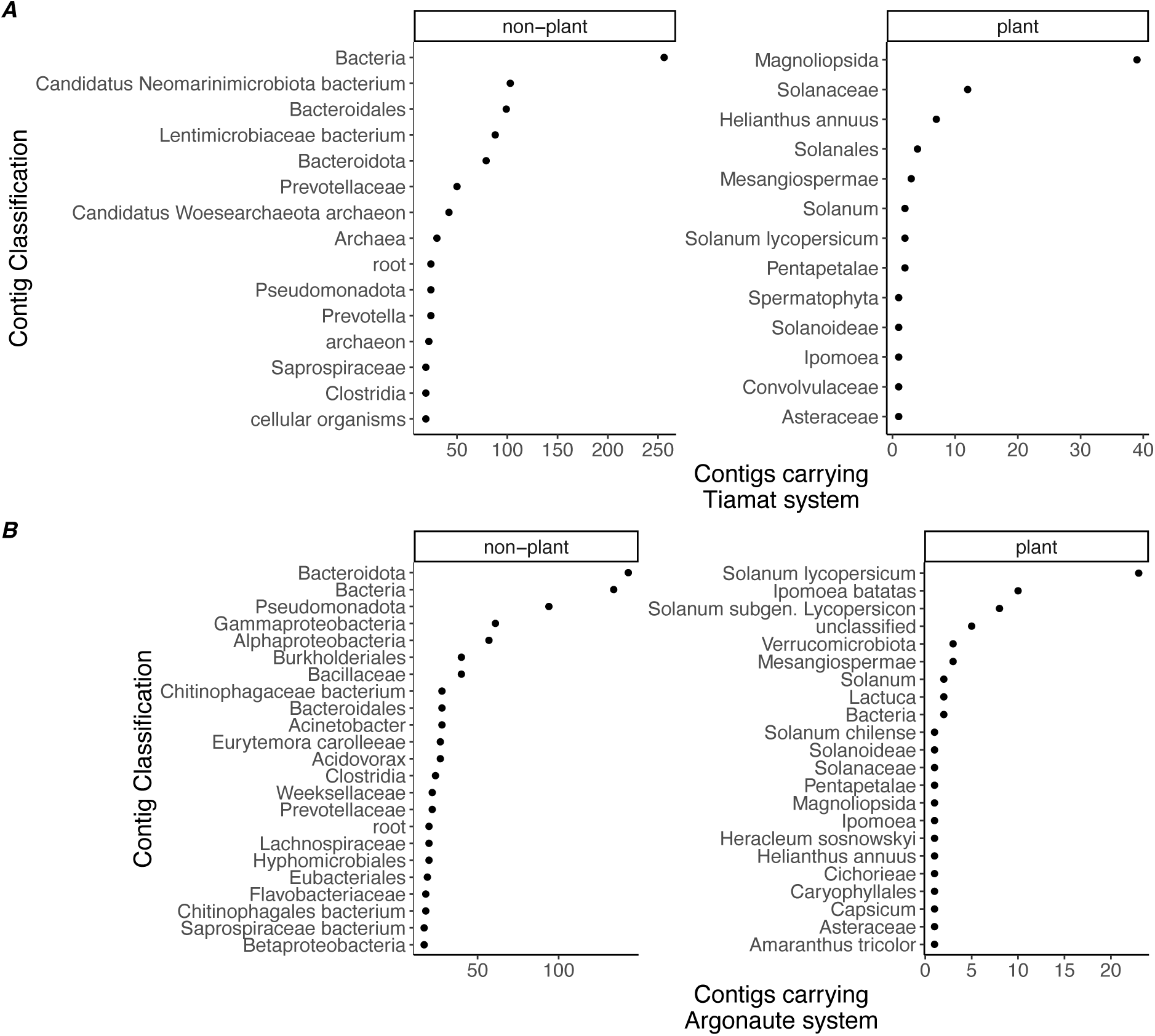
Classifications of contigs carrying Tiamat or Argonaute systems. Counts of contigs extracted from the metagenome assemblies that were identified as carrying a Tiamat (A) or Argonaute (B) system. These contigs were classified using MMseqs2 against the NCBI nr nucleotide database. Counts of contigs are grouped by each last common ancestor (LCA) prediction. Samples were pooled according to plant or non-plant origin. Data shown encompass all classified contigs from the plant samples, but only the corresponding number of ranked LCAs for the non-plant samples.

**Fig. S7.**
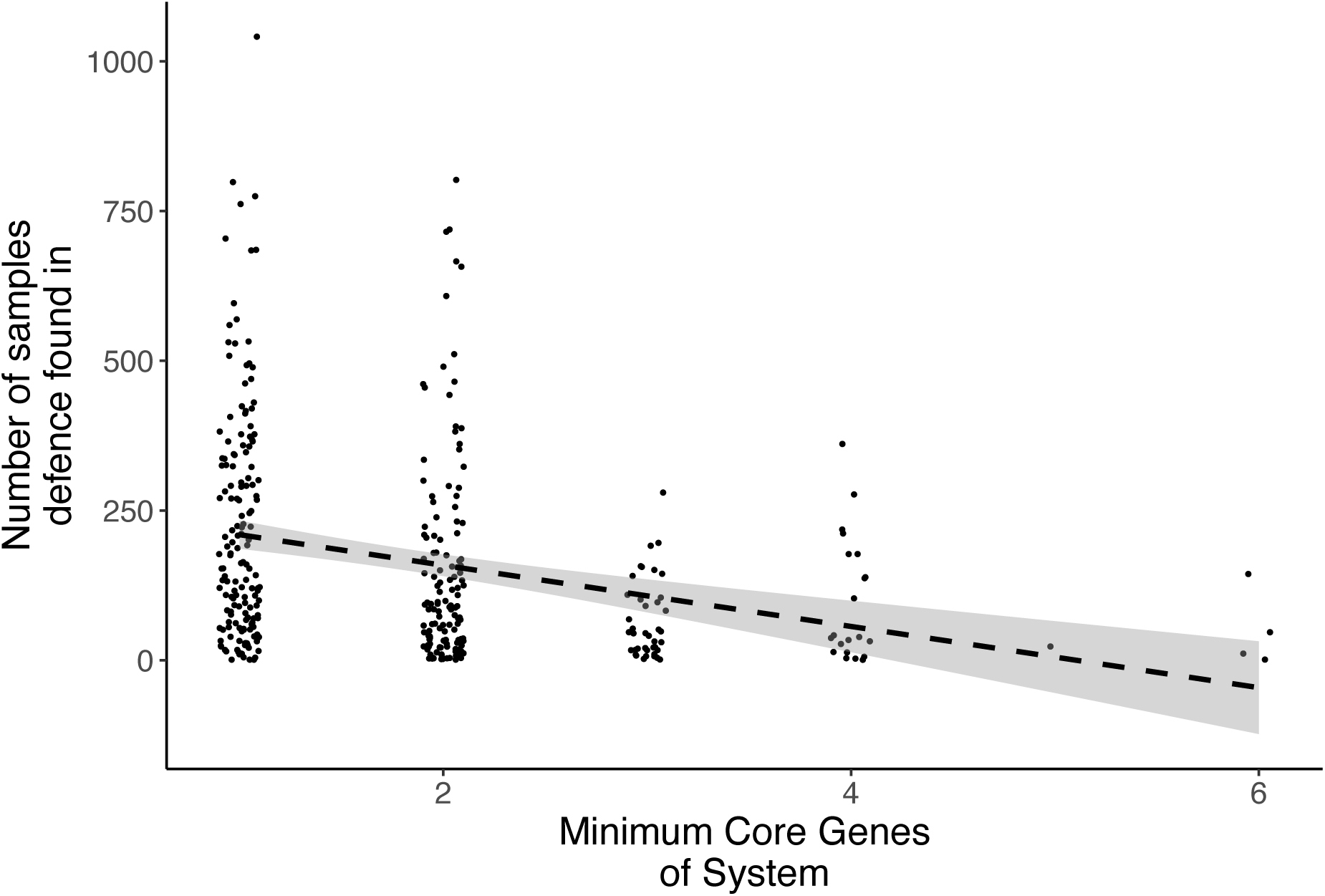
Prevalence of defence system and number of minimum core genes. The minimum number of core genes required for each system was extracted from the PADLOC database system files. These values were plotted against the prevalence of each system i.e. the number of samples in which we detected that defence system. Dashed line represents a linear model fit and shaded areas represent 95% confidence intervals.

## Notes

### Competing Interest Statement

The authors have declared no competing interest.

